# Stabilizing *Plasmodium falciparum* Proteins for Small Molecule Drug Discovery

**DOI:** 10.64898/2026.01.24.701522

**Authors:** Marius Amann, Tim Sträßer, Oliver Einsle, Stefan Günther

## Abstract

Early-stage drug discovery relies on the availability of stable protein for reliable biophysical characterization of ligand binding. However, many *Plasmodium falciparum* proteins are challenging to produce in heterologous systems, which limits their experimental utility. To address this, we tested whether ProteinMPNN-guided sequence design could generate stabilized surrogate constructs that retain wild-type-like structure and binding thermodynamics. Designs were generated with constraints to maintain conserved and binding-site residues for three therapeutically relevant targets: *Pf*BDP1-BRD, *Pf*BDP4-BRD, and *Pf*K13-KREP. The resulting constructs showed markedly increased thermal stability. Using *Pf*BDP1-BRD as a benchmark, isothermal titration calorimetry confirmed that the stabilized variants retained wild-type-like binding thermodynamics with a known ligand. Extending this approach to other targets, a *Pf*K13-KREP construct led to an apo structure with a binding pocket closely matching the wild type, and a stabilized *Pf*BDP4-BRD surrogate - a previously unstable target - enabled the identification of *Pf*BDP4-BRD binders and a 1.25 Å co-crystal structure with a newly found inhibitor. Our findings demonstrate that computationally stabilized surrogates are practical and effective tools for robust biophysics and structure-enabled drug discovery against otherwise challenging malaria proteins.

## 1. Introduction

Target-based drug discovery for *P. falciparum* is often hindered by poor recombinant protein behavior; many targets misfold, aggregate, or are produced at low levels in standard heterologous hosts (Birkholtz et al. 2008). This instability complicates the foundational experiments of any early-stage campaign, namely measuring binding affinities, and obtaining high-resolution structures to guide optimization. Stabilized surrogate constructs could therefore dramatically accelerate these earliest discovery stages.

Recent advances in deep learning–based structure prediction, exemplified by AlphaFold2, have reshaped structural biology by making near–atomic models widely accessible for a broad range of protein families (Jumper et al. 2021). In parallel, the inverse-folding model ProteinMPNN has recently emerged as a particularly powerful approach for sequence design on fixed protein backbones (Dauparas et al. 2022). This has been widely used to improve expression, stability, and function across diverse structural classes (Goverde et al. 2024; Nikolaev et al. 2024; Sumida et al. 2024; D Sa et al. 2025). Despite these advances, a key question for drug discovery remains: Can stability be engineered without distorting the precise binding pocket geometries and thermodynamic profiles that govern small-molecule recognition?

Here, we address this question by applying a ProteinMPNN-guided design strategy to three challenging *Plasmodium falciparum* targets: the bromodomains of *Pf*BDP1 (PF3D7_1033700) and *Pf*BDP4 (PF3D7_1475600), which play a vital role in parasite gene regulation; and the *Pf*K13 Kelch-repeat propeller domain (*Pf*K13-KREP; PF13_0238), associated with artemisinin resistance and parasitic survival (Coppée et al. 2019; Ali et al. 2021; Quinn et al. 2022). Our approach incorporated explicit constraints to preserve biological function, retaining evolutionarily conserved residues (identified by ConSurf) and those within a 12 Å shell for *Pf*BDP1-BRD and *Pf*BDP4-BRD around the known or putative binding pockets (Ashkenazy et al. 2016). For *Pf*K13-KREP, an overlay with the human homolog (PDB 3ZGC) was used to define the corresponding pocket, and residues within an 8 Å shell around this region were preserved. By assessing the resulting surrogates, we evaluated their stability and determined their overall suitability for inclusion in a small-molecule drug discovery pipeline based on their structural integrity and experimental behavior.

## 2. Results

### 2.1 ProteinMPNN yields stable variants while preserving functional sites

We generated potential variants for *Pf*BDP1-BRD, *Pf*BDP4-BRD, and *Pf*K13-KREP using their experimentally determined structures as backbones. For each backbone, we constrained residues selected by evolutionary conservation using ConSurf (conservation score ≥ 6), proximity to the ligand-binding pocket, and structural flexibility (Ashkenazy et al. 2016). The resulting sequences were repredicted using AlphaFold2 and then assessed for structural fidelity, using AlphaFold2 confidence metrics (pLDDT), backbone similarity to the starting structure quantified by TM-score (US-align), solvent-accessible surface area (SASA), and solubility predictors (NetSolP) (Jumper et al. 2021; Thumuluri et al. 2022; Zhang et al. 2022). Constructs that met the criteria were then prioritized, and the most sequence-diverse candidates were selected for experimental validation.

### 2.2 Target-dependent changes in solubility and surface hydrophobicity

As summarized in Table I, NetSolP predictions indicated improved solubility for all representative redesigned constructs compared with their corresponding wild types, while usability (a combined metric integrating solubility and expressibility) improved most clearly for the bromodomains. For *Pf*BDP1-BRD and *Pf*BDP4-BRD, usability increased above 0.5 in the representative designs (*Pf*BDP1-BRD: 0.377 to 0.515; *Pf*BDP4-BRD: 0.383 to 0.537), consistent with improved tractability. In contrast, *Pf*K13-KREP showed increased predicted solubility (0.320 to 0.519), but usability remained below 0.5 (0.328 to 0.365). Across redesigned constructs, mean pLDDT values were uniformly high (96.63 to 99.30) and did not, by themselves, distinguish constructs that yielded well-behaved protein from those that did not.

**Table I:** List of protein constructs with associated metrics during the prediction and selection process as well as the experimental inflection temperature (T_i_).

| Protein | *Pf*K13-KREP<sub>WT</sub> | *Pf*K13-KREP<sub>MPNN</sub> | *Pf*K13-KREP-2<sub>MPNN</sub> | *Pf*BDP1-BRD<sub>WT</sub> | *Pf*BDP1-BRD<sub>MPNN</sub> | *Pf*BDP4-BRD<sub>WT</sub> | *Pf*BDP4-BRD<sub>MPNN</sub> |
| --- | --- | --- | --- | --- | --- | --- | --- |
| mean pLDDT | - | 99.03 | 99.30 | - | 96.63 | - | 97.55 |
| Solubility Usability (NetSolP) | 0.320/0.328 | 0.519/0.365 | 0.508/0.361 | 0.666/0.377 | 0.818/0.515 | 0.683/0.383 | 0.796/0.537 |
| Total SASA (Å<sup>2</sup>) | 12017.3 | 12482.6 | 12142.7 | 7327.6 | 7758.2 | 7464.5 | 7539.1 |
| Hydrophobic SASA (Å<sup>2</sup>) | 4055.83 | 2980.3 | 3621.45 | 2358.1 | 1511.3 | 2536.5 | 2280.7 |
| Hydrophobic fraction | 0.34 | 0.24 | 0.30 | 0.31 | 0.20 | 0.30 | 0.25 |
| Theoretical pI | 5.08 | 5.36 | 4.79 | 8.89 | 5.94 | 7.91 | 6.41 |
| T<sub>i</sub> (°C) | - | 88.4 ± 0.2 | - | 43 ± 0.1 | 77.3 ± 0.8 | - | 64.6 ± 0.5 |

Total SASA changed more modestly across redesigned targets (*Pf*K13-KREP: 12,017.3 to 12,482.6 Å^2^ for *Pf*K13-KREP_MPNN_; *Pf*BDP1-BRD: 7,327.6 to 7,758.2 Å^2^; *Pf*BDP4-BRD: 7,464.5 to 7,539.1 Å^2^). Hydrophobic SASA decreased for each redesigned target (*Pf*K13-KREP: 4,055.8 to 2,980.3 Å^2^ for *Pf*K13-KREP_MPNN_; *Pf*BDP1-BRD: 2,358.1 to 1,511.3 Å^2^; *Pf*BDP4-BRD: 2,536.5 to 2,280.7 Å^2^). Hydrophobic surface fraction also decreased across redesigned targets (*Pf*K13-KREP: 0.34 to 0.24 for *Pf*K13-KREP_MPNN_; *Pf*BDP1-BRD: 0.31 to 0.20; *Pf*BDP4-BRD: 0.30 to 0.25), consistent with reduced exposed hydrophobic character and indicating that the reduced hydrophobic SASA reflects a shift in surface composition.

Within the *Pf*K13-KREP series, the construct that did not yield usable protein (*Pf*K13-KREP-2_MPNN_) retained higher hydrophobic exposure (hydrophobic SASA 3,621.5 Å^2^; hydrophobic fraction 0.30) than the working *Pf*K13-KREP_MPNN_ construct (2,980.3 Å^2^; 0.24) (Table I).

Theoretical pI shifted strongly for the bromodomains (*Pf*BDP1-BRD: 8.89 to 5.94; *Pf*BDP4-BRD: 7.91 to 6.41), consistent with the enrichment of mutations toward charged residues (Table I; Figure S2, Supplementary Information), whereas the *Pf*K13-KREP constructs remained near pI ∼5 with only modest shifts (WT 5.08; *Pf*K13-KREP_MPNN_ 5.36; *Pf*K13-KREP-2_MPNN_ 4.79) (Table I).

### 2.3 Stabilized *Pf*BDP1-BRD surrogate preserves wild-type binding thermodynamics

*Pf*BDP1-BRD served as our benchmark, as we had previously characterized its structure and ligand interactions (Amann et al. 2025). Since the ZA-loop is flexible and vital for bromodomain binding dynamics, it was left unchanged in the design. The stabilized variant showed a substantial increase in thermal stability by nanoDSF (ΔT_i_ = +34.3 °C) (Table I). Isothermal titration calorimetry (Figure 2) showed that the design retains WT-like ligand binding with 1:1 stoichiometry (n = 1) and a similar low-micromolar affinity for RMM23 (design K_D_ = 1.56 µM; wild type K_D_ = 1.24 µM).

**Figure 1:**
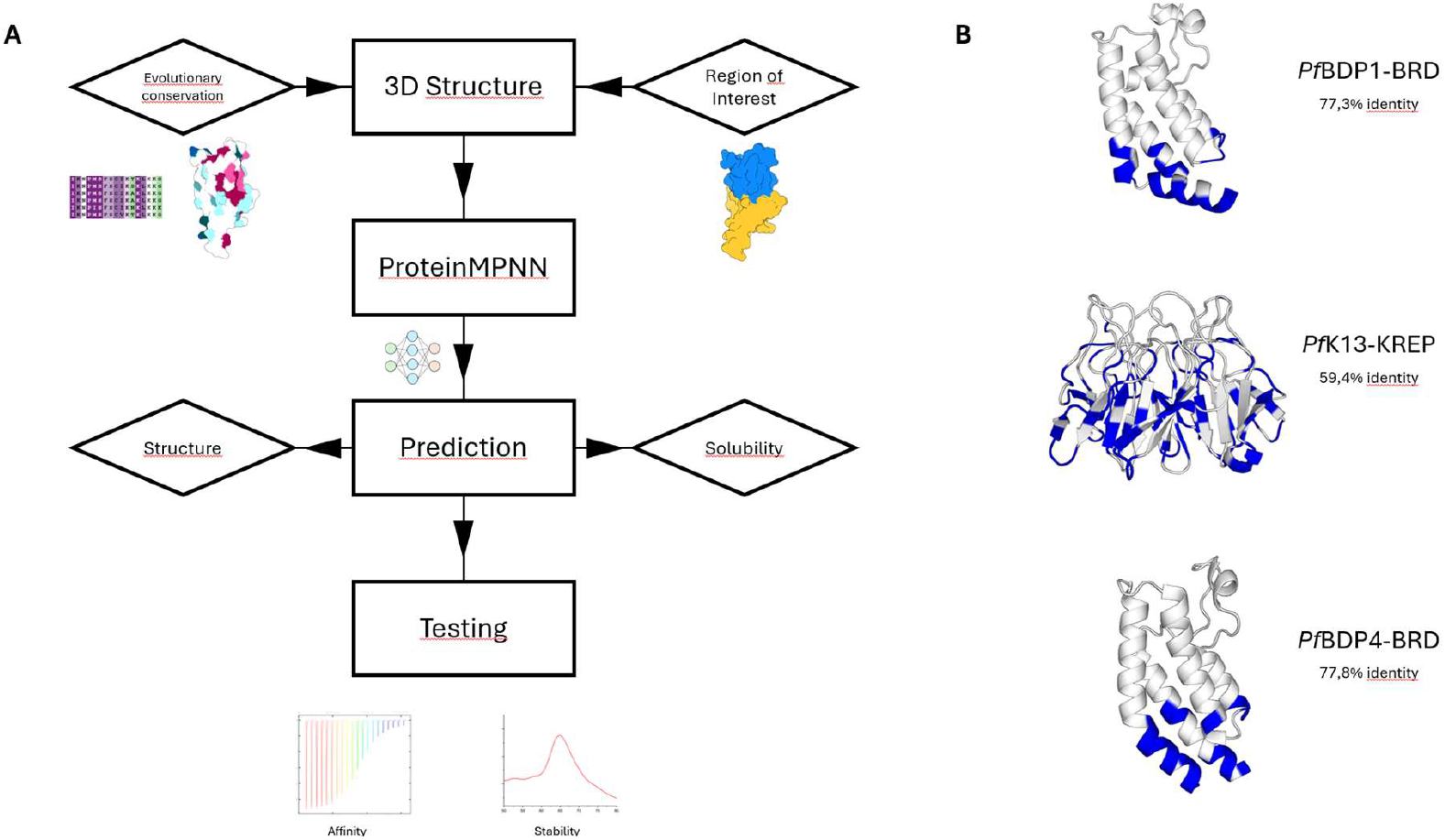
**(A)** Experimentally determined structures were used as fixed backbones. Fixed positions were defined based on evolutionary conservation (ConSurf) and proximity to the ligand-binding pocket to preserve fold and ligand recognition, while remaining positions were redesigned with ProteinMPNN. Designed sequences were repredicted with AlphaFold2 and filtered using model confidence (pLDDT), structural similarity to the backbone by TM-score (US-align), SASA-based surface checks, and predicted solubility (NetSolP). Variants passing thresholds were ranked, and sequence-diverse candidates were selected for experimental validation of stability and ligand binding. **(B)** Structures of the three protein targets with mutational positions colored in blue as well as the sequence identity compared to wild type.

**Figure 2:**
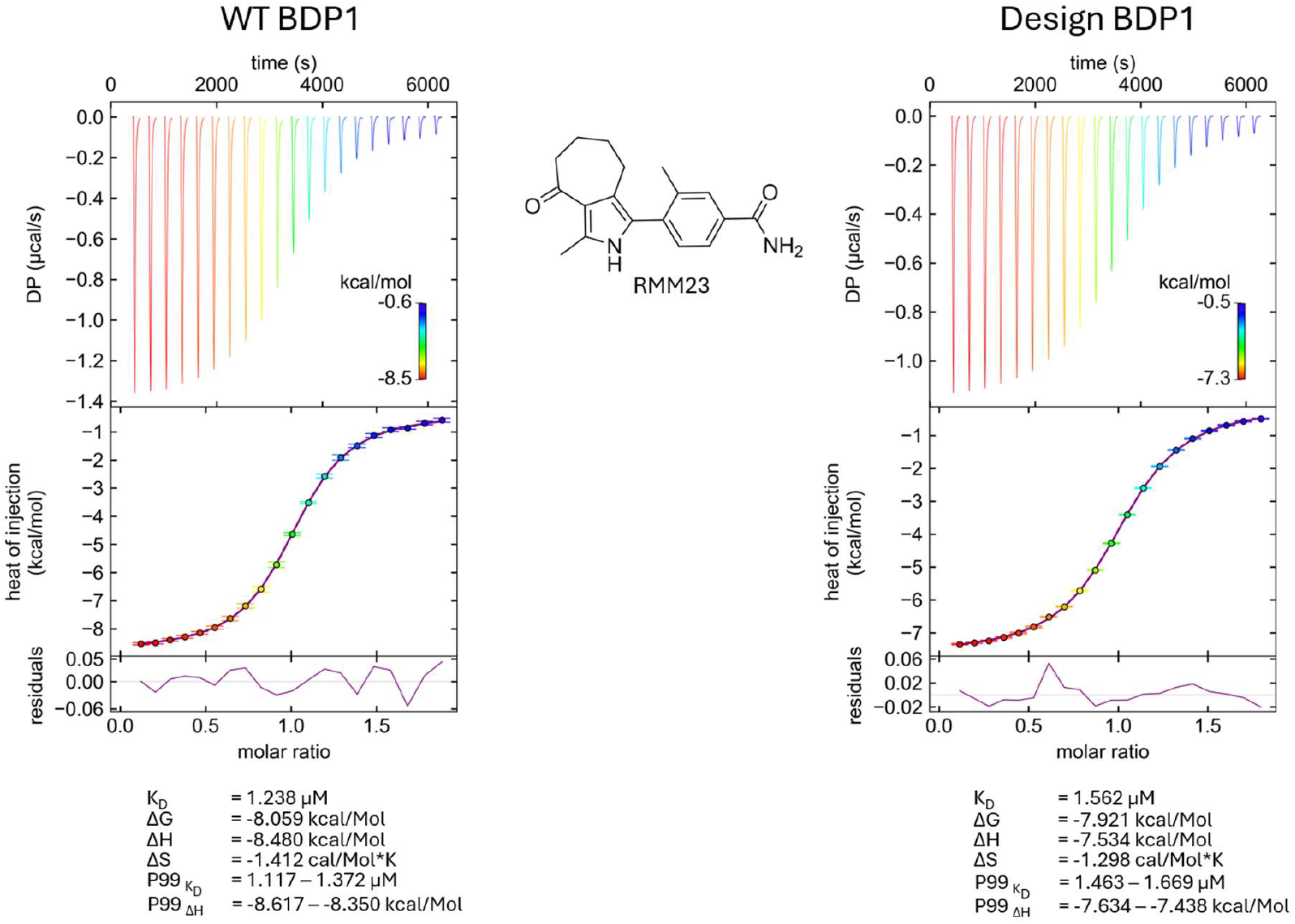
WT *Pf*BDP1-BRD (left) and MPNN stabilized design (right). Top: singular value decomposition (SVD)-reconstructed thermograms. Middle: integrated heats fitted to a one-site binding model, showing a 1:1 stoichiometry (n = 1) and low-micromolar affinity (WT K_D_ = 1.24 µM; design K_D_ = 1.56 µM). Bottom: residual plots of the fits. Dissociation constants and thermodynamic parameters (K_D_, ΔH, −TΔS, ΔG), together with the corresponding P99 intervals, are summarized below each dataset.

### 2.4 Stabilizing PfBDP4 enables structure-based drug discovery

During our experimental preparation, wild-type *Pf*BDP4-BRD was exceptionally unstable, precipitating at room temperature, and therefore unusable for subsequent experiments. In contrast, our stabilized surrogate exhibited a T_i_ of 64.6 °C, rendering it suitable for routine purification and screening. Using these robust constructs, through virtual screening we identified BAY-299 and GSK9311 as probable *Pf*BDP4-BRD binders.

ITC confirmed binding for both, with affinities KD ≈ 101 nM for BAY-299 and ≈ 13 µM for GSK9311 (Figure S1, Supplementary Information). Significantly, a *Pf*BDP4-GSK9311 complex has been solved as a 1.25 Å co-crystal structure. The electron density map unambiguously confirmed a canonical bromodomain binding pose. Overlay with a H4K5ac complexed WT model showed that the local sidechain geometry at the acetyl-lysine binding pocket is similar to the ligand bound form (Figure 3).

**Figure 3:**
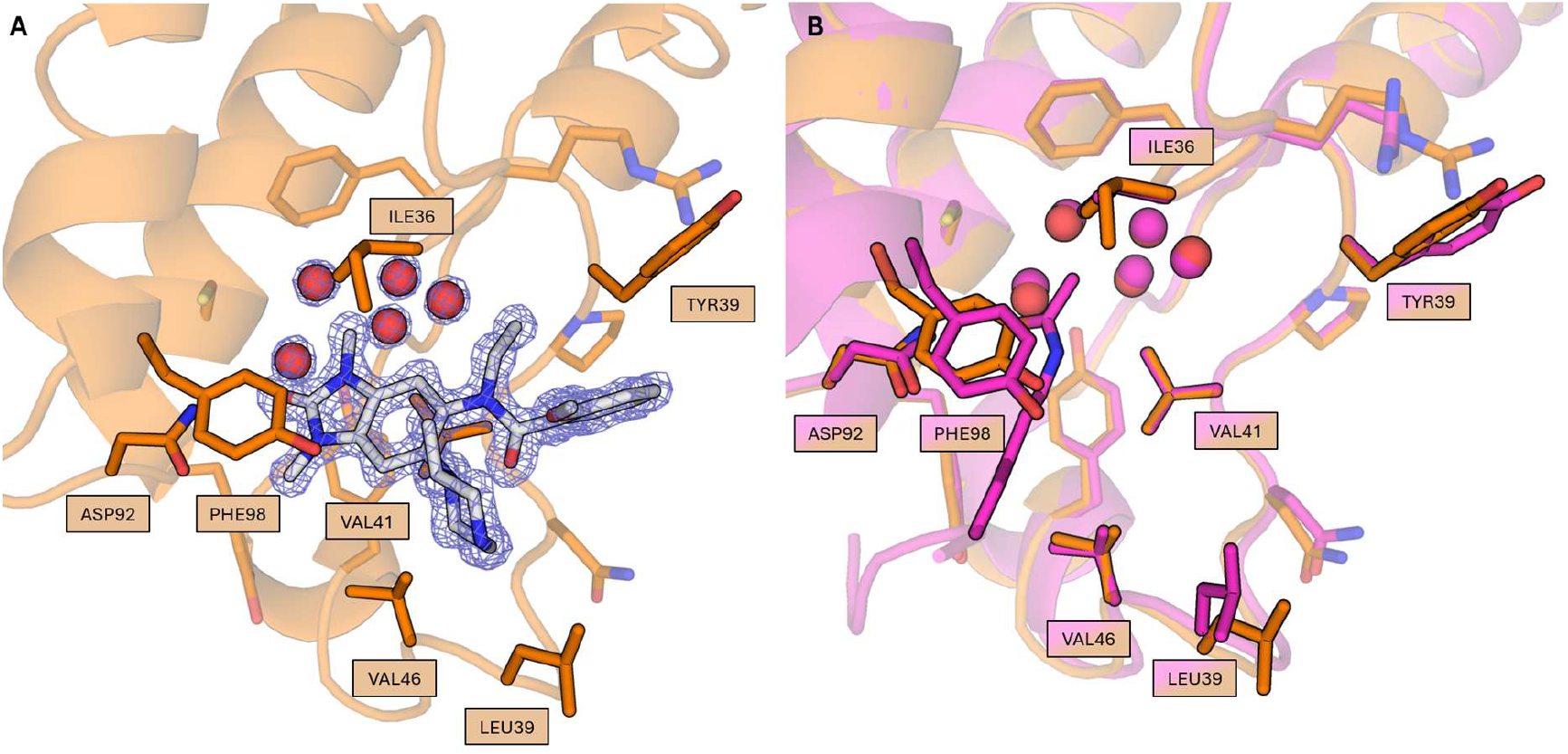
**(A)** *Pf*BDP4-BRD_MPNN_ (orange) with GSK9311 ligand with the 2mFo-DFc composite omit map contoured at 1.5 σ. The benzimidazolone core shows canonical binding to the bromodomain. **(B)** Overlay with H4K5ac complex WT (violet; PDB ID: 5VS7; Cα RMSD: 0.363 Å) shows similar sidechain configuration as with GSK9311.

### 2.5 Stabilized PfK13 propeller yields a 2.15 Å structure with a wild-type-like pocket

Recombinant expression of the *Pf*K13 propeller domain has proven difficult in prior studies (Yan et al. 2022). Our stabilized variant, however, expressed well, crystallized and diffracted to 2.15 Å resolution. Superposition with a wild-type crystal structure (PDB ID: 4YY8) revealed a low backbone RMSD (Cα RMSD: 0.466 Å) and showed that side-chain conformations at the putative shallow binding pocket are fully consistent with the native geometry (Figure 4). Although this was an apo structure, the preserved pocket architecture strongly supports the use of this surrogate for future structure-enabled drug discovery campaigns.

**Figure 4:**
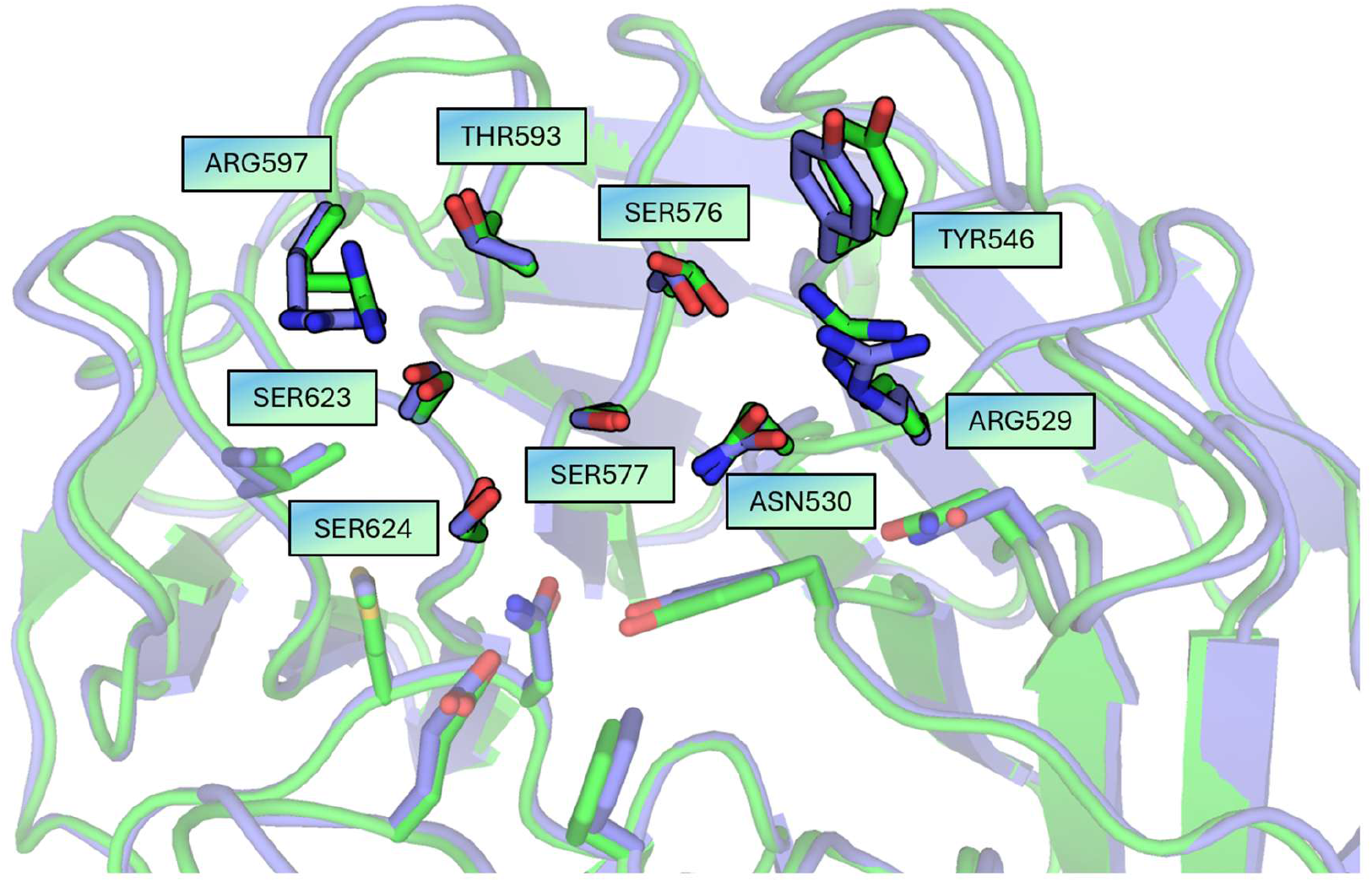
Structural overlay of wild-type (green; PDB ID: 4YY8) vs stabilized PfK13 (blue); close-up of the shallow pocket of the 10% most conserved residues with side chains from wild-type and design overlaid (Cα-RMSD: 0.466 Å).

### 2.6 Crystallization outcomes differ across targets

Stabilization improved biochemical behavior for each target but did not consistently translate into crystallization success. The bromodomain stabilization designs introduced a high fraction of charged substitutions (enriched in Lys/Glu; Figure S2, Supplementary Information). The *Pf*BDP4-BRD surrogate readily formed high-resolution crystals in complex with GSK9311, and the PfK13 surrogate led to an apo structure at 2.15 Å resolution. In contrast, the stabilized *Pf*BDP1-BRD_MPNN_ did not crystallize, despite extensive screening.

Structural inspection of the *Pf*BDP1-BRD wild-type lattice indicates that several mutated sites coincide with crystal-contact regions, suggesting that surface changes introduced during stabilization can alter crystallization propensity in a target-dependent manner (Figure 5).

**Figure 5:**
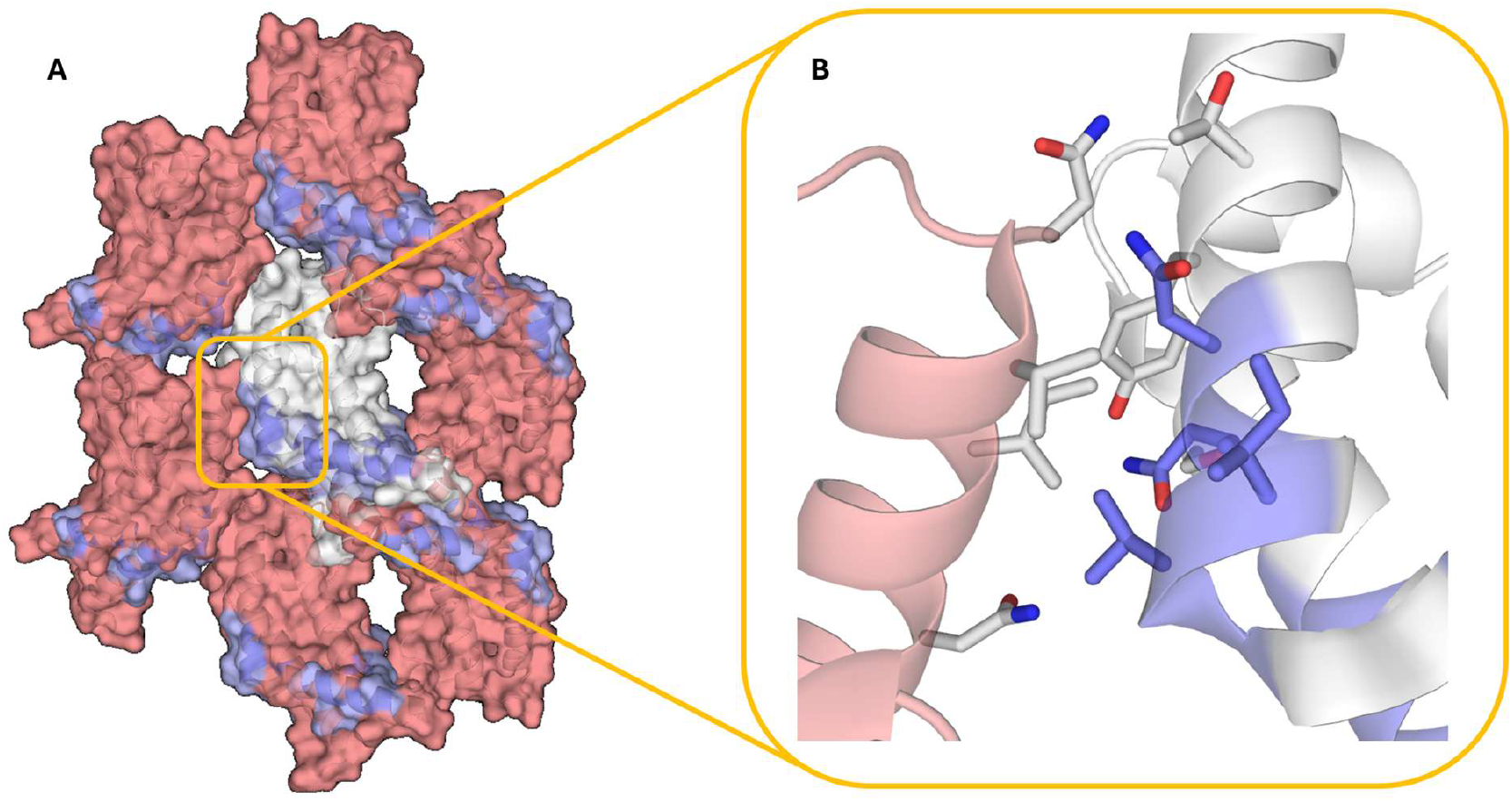
**A:** Crystal structure of *Pf*BDP1-WT (PDB ID: 9HH7), showing the mutated regions in blue; the main structure is shown in white (cartoon) and the symmetry mates in salmon. **B:** Zoomed in view of a crystal contact with mutated residues in blue, non-mutated in white.

## 3. Discussion

Constrained ProteinMPNN redesign produced stabilized versions of three *Plasmodium falciparum* proteins while preserving key ligand-binding features. The redesigned constructs were easier to handle, showed higher thermal stability, and retained the structural elements needed for recognition. This opened experimental routes that were limited with the wild-type proteins, including structure-based drug discovery and biophysical measurements.

A main lesson learned is that no single computational readout was sufficient to predict which designs would work best in the laboratory. AlphaFold2 confidence was high for all redesigns, supporting fold plausibility, but it did not separate designs that later performed well from those that did not. This observation aligns with recent systematic evaluations demonstrating that structure prediction confidence metrics alone do not reliably distinguish experimentally successful protein designs from failures (Garcia et al. 2025). The limitations appear to stem from multiple factors. First, standard AlphaFold2 predictions represent single energy minima and cannot capture the conformational ensembles or dynamic distributions that often govern experimental behavior, particularly for flexible regions and allosteric sites (Karelina et al. 2023; Nussinov et al. 2023). Second, refolding-based validation pipelines show reduced predictive performance when designs remain close to natural sequences, because evolutionary information, most directly the multiple-sequence alignment provided to AlphaFold2, can bias structure prediction toward high-confidence, native-like models even for sequences that later perform poorly experimentally. These metrics also exhibit particular sensitivity to structural flexibility, further complicating their interpretation for designs containing flexible loops or pocket-adjacent regions (Korbeld et al. 2025).

NetSolP scores were more informative, especially for the bromodomains, where the higher-scoring constructs also showed better experimental behavior (Thumuluri et al. 2022). For *Pf*K13-KREP, predicted solubility improved, yet the target still posed greater experimental challenges, consistent with the lower overall score.

Surface analysis provided an important complementary view. Successful redesigns reduced exposed hydrophobic surface area, consistent with a lower tendency to aggregate. Within the *Pf*K13-KREP series, the working construct showed a clearer reduction in exposed hydrophobicity than the non-working variant, which remained closer to wild-type-like hydrophobic SASA and hydrophobic surface fraction (Table I). This suggests that global measures of solvent-exposed hydrophobicity can help flag practical aggregation risk that is not apparent from model confidence metrics alone, and can therefore be assessed alongside more general sequence-based solubility predictors.

In general, a hydrophobic surface character is not necessarily detrimental, particularly when it is functionally required and only transiently solvent exposed. The acetyl-lysine binding site of bromodomains is intrinsically hydrophobic and partly shaped by flexible loops (notably the ZA loop), so loop motion can intermittently expose hydrophobic side chains. Accordingly, some local hydrophobicity can persist near this functional region even as global hydrophobic exposure decreases and predicted scores improve. In such cases, aggregation risk depends strongly on local context: intermittently exposed hydrophobic features in dynamic regions may be less problematic than comparably hydrophobic patches on rigid surfaces that remain continuously solvent accessible.

These observations also point to a limitation of static evaluation. The design and filtering steps rely on a single backbone conformation, but proteins sample conformational ensembles in solution. Flexible loops can expose hydrophobic surfaces only part of the time, which may affect aggregation risk in ways not captured by one structure. Assessing multiple backbone states or adding limited conformational sampling for flexible regions may improve how well computed surface features match experimental behavior.

Against this backdrop, *Pf*BDP1-BRD_MPNN_ provided a useful benchmark for assessing whether stability can be improved without compromising ligand recognition. In this case, the ZA-loop was intentionally left unchanged, as it is flexible and important for bromodomain binding dynamics, while the remainder of the scaffold was redesigned under conservation- and pocket-based constraints. Given the difficulty of predicting which pocket-adjacent mutations will perturb binding-relevant dynamics, we used a conservative constraint strategy preserving residues that directly shape or contact the pocket together with flexible elements likely to influence binding dynamics, while redesigning remaining positions to improve stability. The resulting surrogate showed a strong stabilization by nanoDSF (ΔT_i_ = +34.3 °C) and retained WT-like binding to RMM23 by ITC, with 1:1 stoichiometry and a similar low-micromolar affinity (KD 1.24 µM for WT vs 1.56 µM for the design) (Fig. 2; Table I).

Together, these results support that the constrained redesign strategy can generate a stabilized surrogate that remains suitable for ligand-focused biophysics and early discovery workflows.

Improved stability also did not guarantee crystallization. *Pf*BDP4-BRD_MPNN_ and *Pf*K13-KREP_MPNN_ crystallized and provided high-quality diffraction data; however, *Pf*BDP1-BRD_MPNN_ did not crystallize, despite strong stabilization and good experimental behavior (Table I, Figure 2). *Pf*BDP1-BRD_MPNN_ illustrates that a construct can be an excellent functional surrogate in solution, yet still be a poor crystallization candidate when stabilization mutations remodel lattice-contact surfaces. The bromodomain redesigns introduced many charged substitutions, particularly lysine and glutamic acid residues (Figure S2, Supplementary Information). While such residues are often considered unfavorable for crystallization because they can increase the entropic cost of forming ordered lattice contacts (Derewenda and Vekilov 2006), charge can also support crystallization in a context-dependent manner by enabling directional electrostatic interactions (Nowotny et al. 2019). In our case, *Pf*BDP4-BRD crystallized despite the Lys/Glu enrichment (Figure S2, Supplementary Information), whereas *Pf*BDP1-BRD likely failed because stabilization mutations overlapped with wild-type lattice contacts (Figure 5), consistent with prior observations that surface mutations can disrupt crystal packing, even when they have a stabilizing effect (Hermann et al. 2021).

Importantly, these results underscore that crystallization success remains difficult to predict beforehand and should be treated as a partially independent objective from stability optimization. When structure determination is a central goal, it may be useful to pursue stabilization in parallel with crystallization-oriented construct design, for example by testing additional redesign variants and by avoiding mutations in known or inferred contact-prone surface regions when such information is available.

In summary, this work supports a practical approach for improving difficult malaria proteins for experimental studies: constrain conserved and pocket regions during redesign, then rank candidates using solubility-related predictions together with structure-based surface analysis. The stabilized surrogates and the resulting high-resolution structures show that computational redesign can reduce long-standing experimental barriers and enable ligand-focused studies on proteins that are otherwise hard to work with.

## 4. Materials and Methods

### Computational design workflow

Backbone templates were obtained from the Protein Data Bank: *Pf*K13 Kelch propeller (PDB ID: 4YY8), PfBDP1 bromodomain (PDB ID: 9HH7), and PfBDP4 bromodomain (PDB ID: 4NXJ). Evolutionary conservation was assessed with ConSurf. Residues classified as highly conserved (conservation score ≥ 6) were retained and excluded from designable positions. Consurf MSA parameters were: Maximum number of homologs = 150; Maximum redundancy = 95; MSA building program = MAFFT; Minimum identity = 35; Homolog selection = sample; E-value cutoff = 0.0001

For *Pf*BDP1-BRD and *Pf*BDP4-BRD, the ZA loop was preserved in full to maintain local flexibility and pocket architecture. Ligand-proximal residues were defined as all positions within 12 Å of known or modeled ligands. For *Pf*K13-KREP, pocket-adjacent residues were inferred by superimposing the *Pf*K13 structure (PDB ID: 4YY8) with the human homolog and retaining residues within 8 Å of the homologous site.

All other positions were submitted to ProteinMPNN (weights: v_48_020;Temperature 0.2 and 0.1) for sequence redesign. Cysteine was omitted in sequence redesign. Multiple independent design runs were generated (64 sequences for each Temperature). Candidate variants were repredicted with AlphaFold2 and then filtered via pLDDT ≥ 90 and highest TM-Score to the wild type structure using US-align (Zhang et al. 2022; Yu et al. 2023). Then, sequences were prioritized for experimental testing based on the highest NetSolP solubility and usability scores (≥0.5) as well as solvent-accessible surface area. Final candidates were filtered for sequence diversity among generated candidates using JALview (Waterhouse et al. 2009).

### Surface Property Calculations

Solvent-accessible surface area (SASA) and hydrophobic surface patch metrics were computed using a custom Python workflow implemented as a Google Colab notebook. Per-residue SASA values were computed using FreeSASA (Mitternacht 2016) with default calculation settings (Lee–Richards algorithm; probe radius 1.4 Å) and summed to obtain total SASA. Structures were parsed and prepared in Python using Biopython (Cock et al. 2009) by selecting the analyzed chain and retaining only standard amino-acid residues.

Hydrophobic SASA was defined as the sum of SASA over residues in a specified hydrophobic set (default: C, A, V, I, L, M, F, W, Y, P), and the hydrophobic fraction was calculated as hydrophobic SASA divided by total SASA. Theoretical isoelectric point (pI) was computed from the resulting single-letter amino-acid sequence using Biopython’s ProtParam implementation.

### Virtual Screening Workflow

The crystal structure of *Pf*BDP4-BRD (PF3D7_1475600; PDB ID: 5VS7) was prepared with the Protein Preparation Workflow implemented in Schrödinger Release 2022-3 (Schrödinger, LLC, New York, NY, USA), using default settings. During the energy minimization step, only hydrogen atoms were relaxed using the OPLS4 force field. In addition, five crystallographic water molecules considered important for ligand binding were retained in the binding pocket.

SMILES representations of bromodomain-targeting compounds (9271 molecules) were obtained from ChEMBL. These ligands were processed with LigPrep (Schrödinger Release 2022-3, Schrödinger, LLC) and subsequently docked into the *Pf*BDP4-BRD binding site using the SP (standard precision) mode in Glide (Schrödinger Release 2022-3, Schrödinger, LLC). Docking scores were calculated following post-docking minimization. Top-ranked candidates were visually inspected; selected compounds were purchased from a commercial supplier (MedChemExpress).

### Plasmid construction

Plasmids encoding designed variants were derived from Baker Lab plasmid LM627 (Wicky et al. 2022) and a modified pNIC28-bsa4 vector. LM627 was a gift from David Baker and obtained through Addgene (Addgene plasmid #191551; RRID:Addgene_191551). LM627 already contains a ccdB negative-selection cassette. For pNIC28-bsa4, a ccdB cassette was introduced at the cloning site by FastCloning (Li et al. 2011) or Gibson Assembly (Gibson et al. 2009). Overlapping DNA fragments with 15–25 bp homologous ends were PCR-amplified. For FastCloning, DpnI-treated PCR products with plasmid-matching overhangs were directly transformed into chemically competent ccdB-tolerant E. coli for in vivo assembly. Alternatively, PCR products were purified and subjected to Gibson Assembly prior to transformation into ccdB-tolerant E. coli. Designed variants were assembled by Golden Gate assembly (Engler et al. 2008). All plasmid backbones contained ccdB at the insertion site, which was replaced by the gene of interest during Golden Gate cloning; transformations were performed in ccdB-sensitive E. coli DH5α cells. Following a 1 h outgrowth at 37 °C, cells were plated on agar containing kanamycin. Positive clones were verified via Sanger sequencing using T7 primers.

### Cell growth

Baffled 2 L Erlenmeyer flasks were prepared, each containing 1 L of autoclaved TB medium. Kanamycin (50 μg/L) was added, and the medium was inoculated with 10 mL of preculture. Cells were incubated at 37°C and 180 rpm, while growth was monitored by measuring the optical density at 600 nm. Gene expression was induced at an OD600 of ∼2 by adding IPTG (1 mM for *Pf*K13-KREP variants and 0.2 mM for *Pf*BDP1-BRD_MPNN_ and *Pf*BDP4-BRD_MPNN_). Following induction, cultures were incubated overnight at 16°C and 180 rpm, then harvested at 5000 × g for 12 min using a JLA 8.1000 rotor in an Avanti J-26 XP centrifuge (Beckman Coulter).

### Cell lysis

Cell pellets were resuspended at 3× v/w in cold lysis buffer and stirred until homogeneous. Cells were lysed using a Branson Digital Sonifier (Emerson) with the following settings: 3 s pulse / 7 s pause at 70% amplitude, continuing until a total pulse time of 10 min was reached. The lysates were clarified by centrifugation at 80,000 × g for 1 h at 4 °C.

### Purification (HisTrap affinity and SEC)

His-tagged proteins were purified on HisTrap HP columns (5 mL; Cytiva/GE Healthcare) equilibrated in lysis buffer using an ÄKTA Pure or Prime system. Clarified lysate was loaded at 1 mL.min^−1^, followed by a wash of 10 column volumes of lysis buffer. Proteins were eluted with a 10–300 mM imidazole gradient in the elution buffer. Fractions containing the target protein were pooled, analyzed by SDS–PAGE, and concentrated using centrifugal concentrators (MWCO 10 kDa for *Pf*BDP1-BRD_MPNN_ and *Pf*BDP4-BRD_MPNN_; 30 kDa for *Pf*K13-KREP_MPNN_; Vivaspin 20, Sartorius) at 4,800 rpm, 4 °C. Final polishing was done by size-exclusion chromatography (SEC) into the buffers listed below.

Lysis Buffer for HisTrap: 10 mM Imidazole, 20 mM HEPES pH 7.5, 200 mM NaCl, Glycerol 5% (v/v)

Elution Buffer for HisTrap: 300 mM Imidazole, 20 mM HEPES pH 7.5, 200 mM NaCl, Glycerol 5% (v/v)

SEC buffer (*Pf*K13-KREP_MPNN_): 20 mM HEPES pH 7.5, 100 mM NaCl.

SEC buffer (*Pf*BDP1-BRD_MPNN_ and *Pf*BDP4-BRD_MPNN_): PBS (pH 7.4; 137 mM NaCl, 2.68 mM KCl, 10 mM Na_2_HPO_4_, 1.98 mM KH_2_PO_4_).

### nanoDSF thermal stability assays

Thermal stability was measured by nanoDSF on a Tycho NT.6 (NanoTemper Technologies). Ten microlitres of each sample (100 μM) in SEC buffer were loaded into capillaries and scanned from 35 to 95 °C. Inflection temperatures (T_i_) were calculated with Tycho NT.6 software v1.2.0.750 from the intrinsic fluorescence ratio (350/330 nm). Measurements were performed in technical triplicates.

### Isothermal titration calorimetry (ITC)

Affinity and binding thermodynamics were measured on a MicroCal VP-ITC (Malvern) at 25 °C in PBS. Ligands were prepared as DMSO stock solutions. To exclude heat effects arising from the solvent, a DMSO control was done under the same experimental conditions by adding DMSO to the corresponding buffer (0.04% v/v; ∼5.6 mM) and measuring the protein at the relevant concentration. Since the DMSO control signal was within baseline noise, it was not subtracted from the binding isotherms, to avoid propagating additional variance in the fitted parameters. ITC experiments used ligand in the cell (0.04 mM) and protein in the syringe (0.4 mM). For BAY-299 and *Pf*BDP4-BRD_MPNN_, concentrations were reduced to 0.02 mM ligand (cell) and 0.2 mM protein (syringe). A stirring speed of 351 rpm was used. After one initial 6 µL priming injection, 20 injections of 13.5 µL were delivered at 5 min intervals. All solutions were degassed prior to usage. The first injection was discarded. Raw thermograms were baseline-corrected and integrated with NITPIC (Scheuermann and Brautigam 2015); global fitting was performed in SEDPHAT (Houtman et al. 2007; Zhao et al. 2015) using a one-site (A+B↔AB) model. Confidence intervals were obtained by Automatic Confidence Interval Search with Projection at a confidence level of 0.99 (P99) with Simplex optimization followed by Marquardt–Levenberg refinement at each step. Plots were prepared with GUSSI (Brautigam 2015).

### Protein crystallization and structure determination

All ligands were solved in DMSO (100 mM stock solution) and mixed with the protein in 5x molar excess. Crystallization was done by sitting-drop vapor diffusion in Intelli-Plate 96-3 low-profile plates (Hampton Research). Drops were dispensed with an OryxNano (Douglas Instruments Ltd.). Initial screening used Crystal Screen HT Index (Hampton). For each condition, two drop ratios were set up with a 600 nL total drop volume: 33% protein and 50% protein (protein at 10 - 20 mg.mL^−1^ in SEC buffer). Plates were incubated at 277 K and 293 K and inspected regularly. Suitable crystals were harvested with nylon loops, briefly transferred into reservoir solution supplemented with 10% (R,R)-2,3-butanediol as cryoprotectant, and flash-frozen in liquid nitrogen.

### Data collection and processing

Diffraction data were collected at the beamlines ID30B (McCarthy et al. 2018) and ID23-2 (Nanao et al. 2022) of the European Synchrotron Radiation Facility (ESRF) at 100 K. Data was processed via the ESRF autoprocessing pipeline using autoPROC (Vonrhein et al. 2011). Additional processing and merging was done with AIMLESS (Evans and Murshudov 2013). Phases were obtained by molecular replacement in Phaser (McCoy et al. 2007) using an appropriate search model for PfK13-KREP_MPNN_ (PDB: 4YY8) and an AlphaFold2-based prediction for PfBDP4-BRD_MPNN_ (Jumper et al. 2021; Mirdita et al. 2022). Automated refinement was carried out in PHENIX (Adams et al. 2010), followed by iterative manual model building and inspection in Coot (Emsley and Cowtan 2004). Final model quality was assessed using MolProbity (Williams et al. 2018). All structure illustrations were prepared with PyMOL (version 3.0.3; Schrödinger, LLC 2015). Structural superpositions with root-mean-square deviation (RMSD) calculations were computed using US-align (Zhang et al. 2022).

## Supplementary Materials

**Fig. S1.**
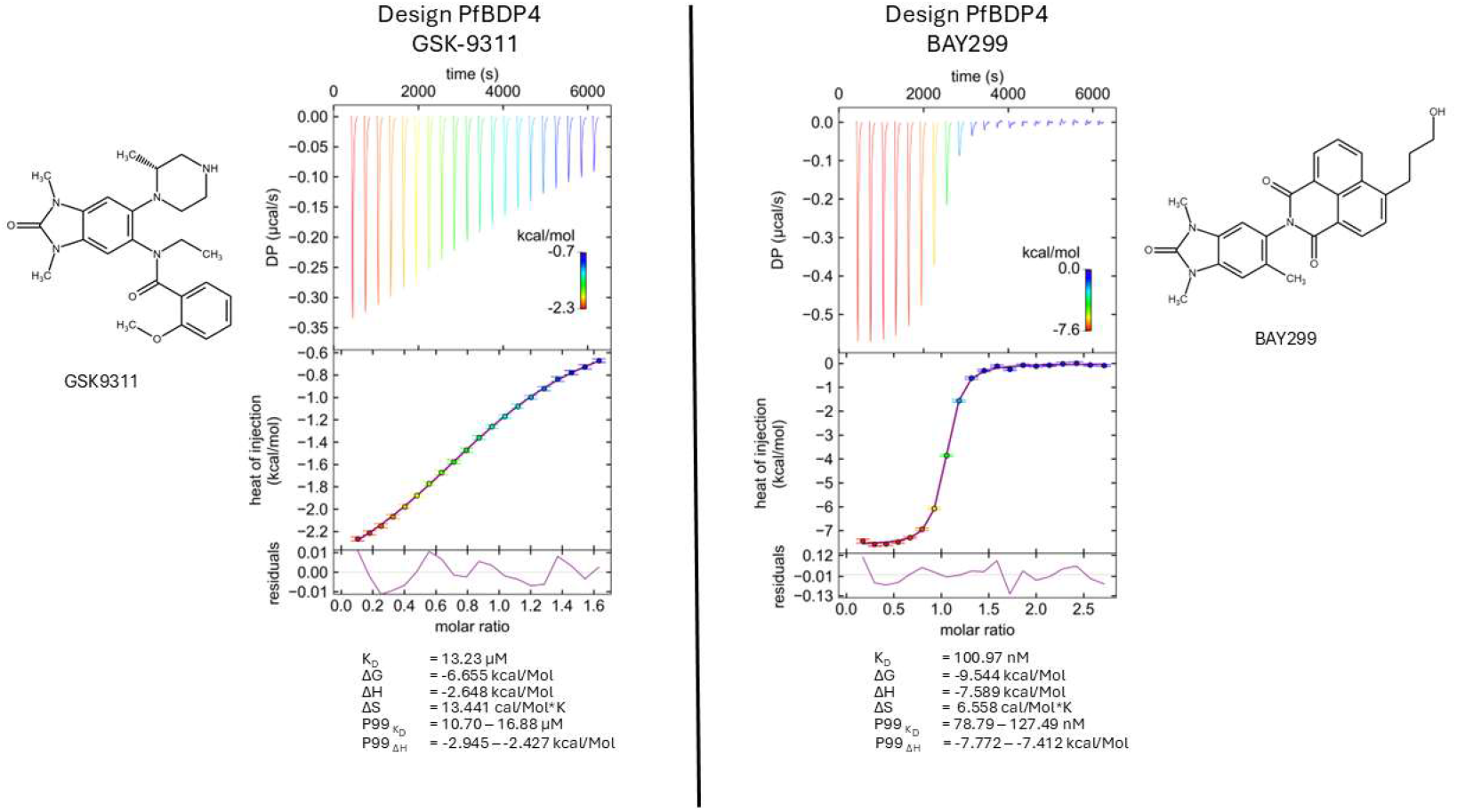
ITC binding of designed PfBDP4-BRD to bromodomain inhibitors. Isothermal titration calorimetry (ITC) of the designed PfBDP4-BRD titrated with GSK9311 (left; K_D_ = 13.23 µM) and (right; BAY-299; K_D_ = 100.97 nM). Top: singular value decomposition (SVD)-reconstructed thermograms. Middle: integrated heats fitted to a one-site binding model. Bottom: residual plots of the fits. Dissociation constants and thermodynamic parameters (K_D_, ΔH, −TΔS, ΔG), together with the corresponding P99 intervals, are summarized below each dataset.

**Fig. S2.**
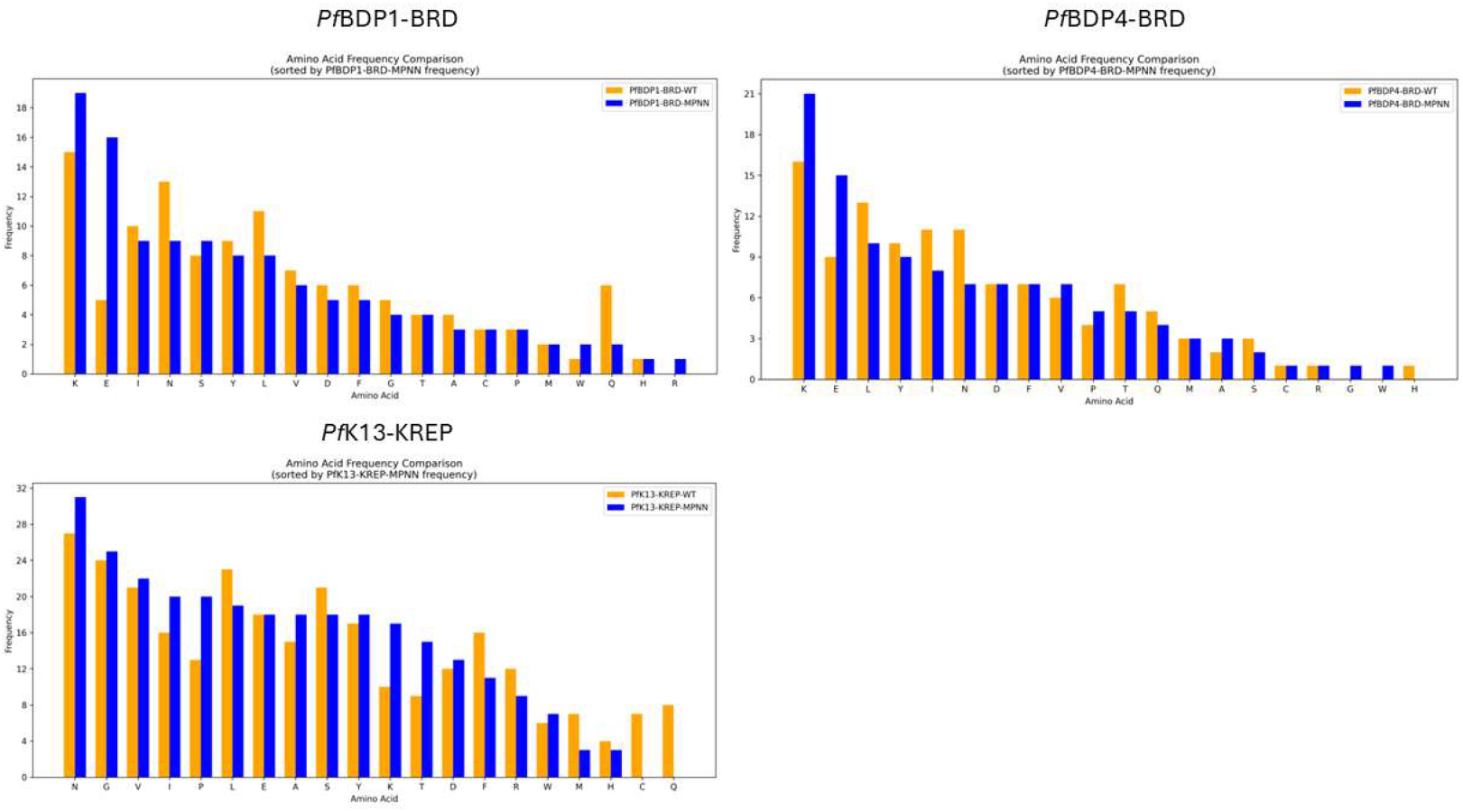
Residue Distribution in ProteinMPNN-Generated Mutants. The frequency of each amino acid substitution is shown, with mutant residues colored blue and wild-type residues in orange. Three protein domains are displayed: *Pf*BDP1-BRD (top panel), *Pf*BDP4-BRD (right panel), and *Pf*K13-KREP (bottom panel). ProteinMPNN-generated mutations show a marked enrichment for charged residues, particularly lysine and glutamic acid, in *Pf*BDP1-BRD and *PfB*DP4-BRD.

**Supporting Table SI.** X-ray data collection and refinement statistics.

|  | **PfK13-KREP<sub>MPNN</sub>** | **PfBDP4-BRD<sub>MPNN</sub> - GSK9311** |
| --- | --- | --- |
| **Wavelength [Å]** | 0.87313 | 0.8731 |
| **Resolution [Å]** | 90.53 - 2.15 | 59.12 - 1.25 |
| **(high resolution shell)** | 2.21 - 2.15 | 1.27 - 1.25 |
| **Space group** | P 31 | P 1 21 1 |
| **Unit cell (a, b, c)** | 92.440 92.440 90.544 | 37.408 32.153 60.426 |
| **(α, β, γ)** | 90.0 90.0 120.0 | 90 101.921 90 |
| **Total reflections** | 357682 (30017) | 232566 (8792) |
| **Unique reflections** | 47058 (3866) | 38945 (1813) |
| **Multiplicity** | 7.6 (7.8) | 6.0 (4.8) |
| **Completeness (%)** | 100.0 (100.0) | 99.3 (94.9) |
| **Mean I/σ (I)** | 8.6 (1.6) | 20.8 (5.5) |
| **Wilson B-factor** | 34.67 | 10.68 |
| **Rmerge** | 0.166 (1.247) | 0.048 (0.257) |
| **Rmeas** | 0.178 (1.335) | 0.052 (0.288) |
| **Rpim** | 0.064 (0.475) | 0.021 (0.128) |
| **CC1/2** | 0.996 (0.677) | 0.999 (0.949) |
| **Number of non-hydrogen atoms** | 7129 | 1394 |
| **Macromolecules** | 6736 | 1023 |
| **Ligands** | 18 | 42 |
| **Waters** | 375 | 329 |
| **RMS (bonds)** | 0.003 | 0.008 |
| **RMS (angles)** | 0.65 | 0.90 |
| **Ramachandran favored (%)** | 97.66 | 100.00 |
| **Ramachandran allowed (%)** | 2.34 | 0.00 |
| **Ramachandran outliers (%)** | 0.00 | 0.00 |
| **Average B-factor** | 37.45 | 18.31 |

## Data and Code Availability

Data that support the findings of this study are available in the supplementary material of this article. Coordinate files for *Pf*BDP4-BRD complex with GSK9311 and *Pf*K13-KREP as well as the protein sequences used in this work will be available upon publication of the peer-review article. The Python/Google Colab workflow used to compute surface properties is available on GitHub: https://github.com/mar4mn/Hydrophobic-Surface-Analysis

## Acknowledgments

We would like to thank the European Synchrotron Radiation Facility (ESRF) for their support.

## Author Contributions

**Marius Amann**: Conceptualization (lead); Methodology (lead); Software (lead); Data curation (lead); Formal analysis (lead); Investigation (lead); Validation (lead); Visualization (lead); Writing—original draft (lead); Writing—review & editing (equal).

**Tim Sträßer**: Investigation (lead); Data curation (equal); Validation (equal); Formal analysis (supporting); Methodology (supporting); Writing—review & editing (supporting).

**Oliver Einsle**: Resources (lead); Supervision (equal); Writing—review & editing (supporting)

**Stefan Günther**: Conceptualization (equal); Project administration (lead); Supervision (equal); Writing—original draft (supporting); Writing—review & editing (equal).

